# Experimentally induced pain and paresthesia respond differently to parameter changes of cuff-based compression in pain-free young individuals

**DOI:** 10.1101/2024.07.22.604622

**Authors:** Jacek Skalski, Sylwia Swoboda, Tibor M. Szikszay, Piotr Wodarski, Andrzej Bieniek, Kerstin Luedtke, Wacław M. Adamczyk

## Abstract

Neuropathic pain is a significant therapeutic challenge due to the co-occurrence with other neurological symptoms such as paresthesia. Human-based models such as cuff algometry can enhance our understanding of pain-paresthesia relationships. This experiment aimed to characterize (psychophysically) pain and paresthesia evoked by stimuli of different temporal and intensity parameters and to demonstrate the reliability of experimental induction of these two symptoms using cuff algometry. Forty participants, aged 18-35, were exposed to mechanical pressure stimuli at three intensities (100, 150, 200 mmHg) and three durations (90, 120, 150s). Skin Conductance (SC) was continuously monitored, and participants rated pain and paresthesia in real-time using a computerized visual analog scale. The General Linear Model analysis revealed significant differences in paresthesia across all durations (p<0.01), but not all intensities, as paresthesia did not increase from 150 to 200 mmHg (p>0.05). Conversely, pain responses showed significant differences across all pressure intensities (p<0.05) but not durations, as pain did not increase from 90 to 120 and from 120 to 150s (p>0.05). No interaction effects were found for either symptom. SC analysis showed no significant main or interaction effects. Intraclass correlation coefficients (ICCs) indicated moderate to good reliability for pain and paresthesia induction across different durations and intensities (ICC: 0.52-0.90), while SC showed poor to moderate reliability (ICC: 0.21-0.73). In conclusion, computerized cuff algometry seems to be an effective and reliable method for simultaneously inducing and assessing pain and paresthesia, revealing that these symptoms follow different patterns based on pressure duration and intensity.

## 1. INTRODUCTION

Neuropathic pain (NP) is a common clinical condition with an estimated prevalence of 6.9% - 10% in the general population [50] and may account for as much as 18.1% of chronic pain cases [3]. The effective treatment of NP is a major challenge due to the complexity of the underlying pathophysiology and the limited efficacy of currently available treatments [10]. IASP defines NP as a pain that can arise as a consequence of a lesion affecting the somatosensory system. However, the exact pathophysiological mechanisms, and the interaction between pain and other symptoms that co-exist with NP, are not yet fully understood, highlighting the need for further basic science research.

Neuropathic conditions are often accompanied by positive symptoms such as “paresthesia” [57]. According to the National Institute of Neurological Disorders and Stroke (NINDS) definition, paresthesia refers to a burning or prickling sensation felt in different body regions and occurs during sustained pressure on a nerve. Paresthesia commonly co-occurs with pain in conditions involving nerve compression, such as carpal tunnel syndrome [26,35] and radicular back pain [41] and is also prevalent in disorders such as fibromyalgia, migraine, or in patients with healed burns [4,11,56].

A better understanding of NP requires the simultaneous assessment of pain and paresthesia under controlled conditions, as both symptoms often occur together and influence each other [13,14,49]. However, current studies mostly investigate these sensory disturbances in isolation [19,39,44,46], leading to an incomplete understanding of their interaction. Furthermore, since these two symptoms often co-occur, assessing them in isolation limits the ability to evaluate how an intervention targeting one symptom might influence the other. In contrast, assessing both symptoms simultaneously can reveal whether improvement in one symptom could benefit the other, for instance, through mechanisms such as the placebo effect [55].

A model that elicits both symptoms reliably and in real time, under clinically relevant conditions, could provide valuable insights for research and clinical practice, especially for the development of more precise diagnostic and therapeutic approaches. To the best of our knowledge, no reliability and validity data exist for a technique designated to measure two symptoms simultaneously. However, previously published study results indicate that pressure pain and tolerance thresholds showed good to excellent reliability [18,30]. Whether suprathreshold pain and paresthesia ratings are characterized by good reliability remains to be tested.

Cuff algometry represents a novel approach to the simultaneous induction and assessment of pain and paresthesia through the simulation of nerve compression and ischemia [38,39]. In order to supplement the subjective assessments of pain and paresthesia with objective measures, the recording of skin conductance (SC) is employed, as it provides a physiological proxy of the symptoms, indicating the activity of the autonomic nervous system [16,31].

The objective of this study is to characterize the stimulus-response relationship and assess the reliability of simultaneously inducing and measuring experimental pain, paresthesia and skin conductance in a cuff-based paradigm. In contrast to previous studies, which typically employed separate assessments, this study examines the real-time evolution of pain and paresthesia in response to varying stimulus intensities and durations. Moreover, the assumptions regarding the gradual progression of paresthesia due to hypoxia [20,34,54] and the exponential increase in pain [45] are examined, as are the consistency and reliability of the measurements.

## 2. METHODS

### 2.1. Study overview and general information

The experiment was conducted in the Laboratory of Pain Research at The Academy of Physical Education in Katowice in accordance with the recommendations of the Declaration of Helsinki. Each participant signed an informed consent before taking part in the study. The study protocol was approved by the local bioethics committee (decision no. 9/2019) and registered on the Open Science Framework platform (https://osf.io/wmhtq). All procedures were conducted by the same examiner (S.S).

The study exposed 40 participants to mechanical pressure stimuli using a computerized cuff algometry setup. Stimuli were delivered at varying intensities (100, 150, 200 mmHg) and durations (90, 120, 150s), allowing for the simultaneous assessment of pain and paresthesia. Real-time ratings of both symptoms were collected, and electrodermal activity was continuously monitored throughout the procedure.

### 2.2. Participants and inclusion criteria

A group of 40 participants (20 females, 20 males) were recruited from the university community by using social media advertisement and word-of-mouth. The sample size of 40 was chosen based on practical constraints such as time and resource availability, as well as to satisfy needs for the Central Limit Theorem [2] considering the exploratory nature of the study. This number was considered sufficient to identify initial patterns and generate hypotheses for future larger-scale studies.

Only individuals who declared that they felt subjectively healthy and met the inclusion criteria described below, aged 18-35, were allowed to participate in the study. A screening questionnaire was used to exclude those individuals who had any medical condition that could have influenced pain perception, such as neurological diseases or upper limb pain, lasting more than 24h within the month preceding the study, had any systemic disease or the history of intolerance to ischemic stimuli during routine blood pressure measurement. To standardize the side of the body exposed to ischemic stimuli and to minimize the potential impact of handedness on pain sensitivity, only right-handed participants were included [47,53].

Before signing the informed consent, participants were informed about the study procedures and that they can withdraw from the study at any moment without any consequences and without providing reasons for a withdrawal.

### 2.3. Study design

The study employed a within-subject design. Participants received three intensities of mechanical pressure stimuli (100, 150, 200mmHg) applied with three different durations (90, 120, 150s) in two sessions which resulted in nine stimuli during the first session and nine during the second session (18 stimuli in total). The intensities (100, 150, 200 mmHg) and durations (90, 120, 150s) were selected based on safe pressure levels reported in the literature and duration-related axonal excitability, ensuring effective and safe stimulation [25,38].

Random sequences of parameters were computer generated for each session to reduce potential carry-over and order effects, which could occur if participants experienced the same sequence repeatedly and to prevent habituation, anticipation, or bias in participant responses. During each stimulus, participants rated both symptoms (pain and paresthesia) simultaneously in real time and additionally provided overall ratings after each stimulus.

### 2.4. Experimental procedure

#### 2.4.1. Equipment

To induce pain and paresthesia, a commercially available sphygmomanometer’s cuff (Omron M3 HEM-7200-EE2(V), Omron Healthcare Co., Ltd., Japan) was adapted and paired with sliding potentiometers to rate the symptoms on Computerized Visual Analogue Scales (CoVAS) in real-time. Skin conductance (SC) was measured using electrodermal activity (EDA) and continuously recorded via a USB coupler device (USB Physiology Coupler Contact Precision Instruments®, Pendragon House, Butleigh Rd, UK) connected with two 8mm diameter Ag-AgCl cups filled with EDA gel (EDALYT, EasyCap GmbH, Germany) attached to the distal phalanges of the non-dominant index and middle fingers. The experiment was programmed and carried out using the PsychoPy (v2021.2.3) open-source software [42]. Two Python libraries (“serial” and “u3”) were used to control the cuff, potentiometers, and digital/analogue converter LabJack U3-LV device (LabJack Corporation, Lakewood, CO, USA) which was used to ensure communication with the EDA device.

#### 2.4.2. Preparation

The experiment was conducted in an ISO-certified laboratory with environmental standardization. The room temperature was set to approximately 23°C, and noise was prevented to ensure a quiet, comfortable, and controlled setting for sensory testing. The instruction was standardized and read out by the researcher during the experiment to ensure consistency across participants. The instructions (translated) are provided in **Appendix S8**.

Participants were seated on an armchair next to the computer screen with arms resting on a table. After completing the screening questionnaire and signing the informed consent, age, height, weight and arm circumference (measured with a measuring centimeter at the widest point of the arm, near the mid-length of the humerus) were collected. Additionally, to efficiently assess participants’ fear levels, they were asked to rate their fear of pain on a 0-10 Verbal Rating Scale (VRS) where 0 indicated “no fear of pain” and 10 indicated “very high fear of pain” [1]. A sphygmomanometer cuff was placed on the participant’s left arm, just above the elbow joint (flexed at 90° angle). The phalanges of the index and middle finger were previously prepared by washing the hands with lukewarm water [6]. Two electrodes were attached using self-adhesive tape. The examiner explained how to operate the potentiometers to reflect the intensities of both, pain and paresthesia, before the data collection was started.

#### 2.4.3. Familiarization trial

After the cuff was fitted, one test compression (150mmHg) was performed to familiarize study participants with the modality of the stimuli and the assessment procedure. Participants were asked to maintain a relaxed position during the test, minimize movement of the tested limb (left), and assess the intensity of pain and paresthesia in real time (using their right hand) on the sliders placed in front of them.

#### 2.4.4. Measurement and outcomes

Following the familiarization trial, the main phase of the experiment started. It consisted of two series of nine stimuli each (∼ 120s interval between trials). The stimuli were presented in a random order, unbeknownst to the participants and the examiner, and each parameter was applied twice per participant for a total of 18 compressions to allow for reliability analysis. A 15-minute break between the first and second series included removing the compression cuff and encouraging free limb movement to stimulate blood circulation. The parameters for the interval times were chosen based on the development trials conducted in our lab. During these trials, participants were asked about their sensations following each compression. Before a new stimulus could begin, participants were required to press the space button, which they were instructed to do only when all symptoms had subsided, and the sliders were reset to their starting positions. The 15-minute break between the first and second series was designed to minimize any potential carry-over effects. Removing the compression cuff and encouraging free limb movement during this period helped restore normal blood circulation and ensured that the sensory system was reset to baseline before proceeding with the second series. The second series was identical to the first. Participants assessed pain and paresthesia intensity in real-time using the CoVAS by maneuvering potentiometers. They were instructed as follow: “The slider on the left is used to determine the intensity of the pain you feel, where the bottom indicates no pain, and the top indicates the worst pain you can imagine. The middle slider is used to determine the intensity of the tingling sensation felt, where the bottom indicates no tingling, and the top indicates the worst tingling you can imagine. The slider on the right is inactive and will not be used.” After each trial, they rated overall pain and paresthesia on a VAS scale displayed on the computer screen. Main outcomes included pain and paresthesia intensity for different compression intensities and durations, measured on the CoVAS scale. The EDA signal, including skin conductance (SC), was continuously recorded throughout the experiment, with each stimulus onset and offset digitally marked to capture autonomic nervous system responses associated with pain perception. Additionally, SC was monitored to explore whether it could provide objective traces of emerging paresthesia sensations, as it reflects physiological changes, such as sweat gland activity, driven by sympathetic nervous system activation. Demographic data (sex, age, weight, height) and arm circumference were collected to characterize the sample and assess correlations with pain and paresthesia severity.

### 2.5. Data processing and statistical analysis

Raw data from CoVAS were first imported and preprocessed by using MATLAB R2023b (MathWorks, Natick, MA) prior to the main statistical analyses. Continuously measured pain and paresthesia intensity as well as stimulus data (pressure) were first reduced at the subject level by averaging ratings for each second of data acquisition. Reduced data were then plotted as a time-function and used to calculate mean ratings for each applied stimulus as well as Areas Under the Curve (AUC) for each of the 2 sessions. To correct for the delay in establishing a target level of pressure, parameters (mean, AUC) were calculated based on the data from the plateau phase of each stimulus. The skin conductance (SC) signal was first smoothed with the moving average and later encompassed with similar preprocessing steps as the behavioral data described above. Data from ten participants were excluded from analysis due to missing signal recordings in the first (n = 4) or second (n = 2) session of the experiment or artefacts (n = 4) as the result of device malfunction during data acquisition.

Descriptive statistics are presented as means and standard deviations (SDs) unless otherwise indicated. The main analysis was performed by means of the General Linear Model (GLM) for repeated measures, with two within-subject factors, i.e., stimulus intensity (pressure of 100, 150, 200mmHg) and stimulus duration (90, 120, 150s). Primary analysis was performed on the extracted means from curves instead of AUC to be able to analyze the effect of two factors (time and stimulus intensity) independently due to relative dependency of AUC values from both reported symptoms and duration of exposure. Post-hoc Tukey tests were performed in case of statistically significant main and/or interaction effects. Steven’s power function was fitted to the aggregated stimulus-response function to extract the exponent indicating the growth trajectory (e.g., exponential vs. logarithmic). Furthermore, reliability of measurement was calculated by using Intraclass Correlation Coefficient (ICC_(3,1)_) and Smallest Detectable Differences (SDDs) to reflect the correlation between the data from the first session vs. second session. Reliability was interpreted as excellent (0.9-1), good (0.75-0.9), moderate (0.5-0.75) or poor (<0.5) [28]. Preliminary concurrent validity was analyzed by applying Pearson-product moment correlation coefficient and correlating overall ratings collected after application of the stimulus with corresponding averaged ratings derived from continuously measured symptoms (pain and paresthesia). Bland-Altman plots were used to further visualize systematic differences between the two methods and to explore bias in datasets from the two sessions; 95% Limits of Agreement (LoA) were calculated.

## 3. RESULTS

Descriptive statistics are presented in **Table 1**, whereas means and standard deviations for all experimental conditions and outcomes are presented in **Table 2**. Averaged outcomes measured over-time and stimulus data are shown in **Figure 1**.

**Table 1.** Descriptive statistics.

| **Variable** | **Mean** | **SD** |
| --- | --- | --- |
| *<u>Participant characteristics</u>* |  |  |
| Age [years] | 21.10 | 1.72 |
| Body mass [kg] | 72.06 | 14.36 |
| Height [cm] | 174.15 | 8.69 |
| Arm circumference [cm] | 28.85 | 4.06 |
| Sex [% of females] | 50% |  |
| *<u>Psychological measures</u>* |  |  |
| Fear of pain [VRS] | 3.28 | 1.69 |

**Legend:** Fear of pain was measured on a 0 – 10 Verbal rating Scale (VRS) scale (0 – no fear of pain; 10 - the worst fear of pain imaginable).

**Table 2.**
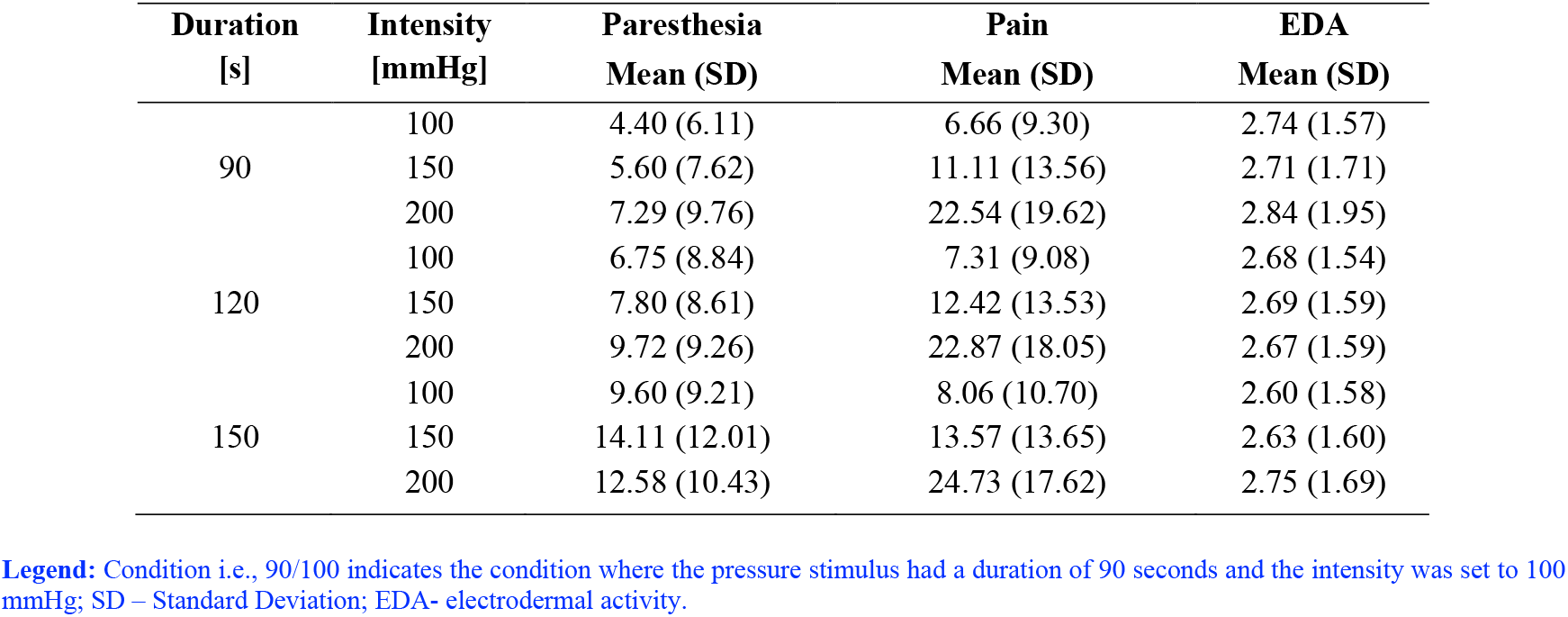
Means and SDs for all conditions and outcomes.

| **Duration**<br>**[s]** | **Intensity**<br>**[mmHg]** | **Paresthesia**<br>**Mean (SD)** | **Pain**<br>**Mean (SD)** | **EDA**<br>**Mean (SD)** |
| --- | --- | --- | --- | --- |
| 90 | 100 | 4.40 (6.11) | 6.66 (9.30) | 2.74 (1.57) |
|  | 150 | 5.60 (7.62) | 11.11 (13.56) | 2.71 (1.71) |
|  | 200 | 7.29 (9.76) | 22.54 (19.62) | 2.84 (1.95) |
| 120 | 100 | 6.75 (8.84) | 7.31 (9.08) | 2.68 (1.54) |
|  | 150 | 7.80 (8.61) | 12.42 (13.53) | 2.69 (1.59) |
|  | 200 | 9.72 (9.26) | 22.87 (18.05) | 2.67 (1.59) |
| 150 | 100 | 9.60 (9.21) | 8.06 (10.70) | 2.60 (1.58) |
|  | 150 | 14.11 (12.01) | 13.57 (13.65) | 2.63 (1.60) |
|  | 200 | 12.58 (10.43) | 24.73 (17.62) | 2.75 (1.69) |

**Legend:** Condition i.e., 90/100 indicates the condition where the pressure stimulus had a duration of 90 seconds and the intensity was set to 100 mmHg; SD – Standard Deviation; EDA- electrodermal activity.

**Figure 1.**
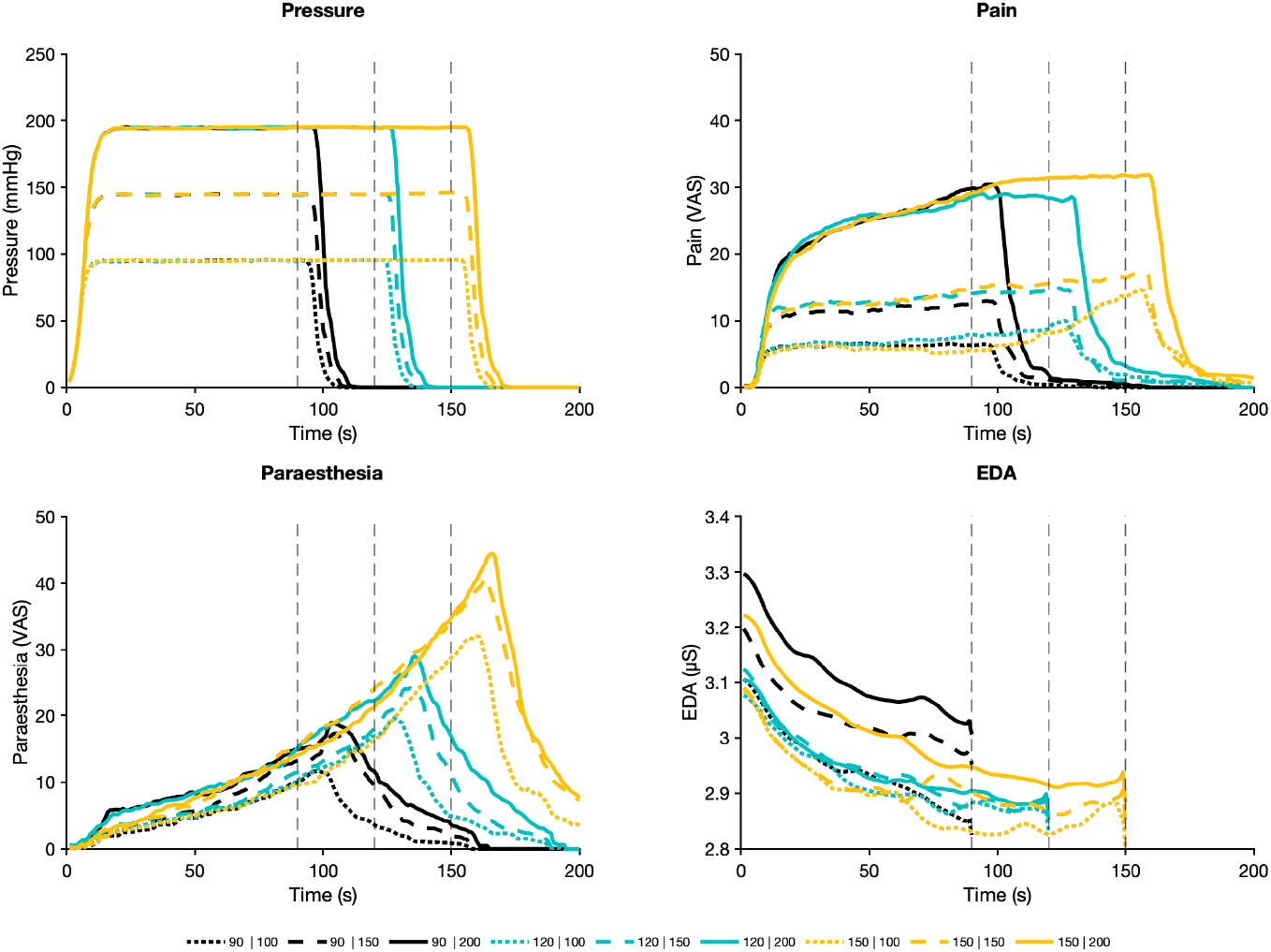
Main outcome measures and stimulus data collected in real time. The upper left panel represents the pressure (mmHg) from each stimulus type. The upper right panel reflects mean pain scores; the lower left mean paresthesia scores, the lower right mean SC values in μS. Each line represents a different stimulus. Colour (blue, cyan, purple) represents stimulus duration (90, 120, 150s.), line type (dotted, dashed, continuous) represents stimulus intensity (100, 150, 200mmHg). Vertical markers indicate stimuli offsets.

Primary analyses reported below were conducted on average perceived pain and paresthesia intensity, however, they were repeated also with AUC as an outcome and are presented separately in the **Appendix S2-S4**.

### 3.1. Paresthesia

The GLM analysis revealed statistically significant main effects for “time” (F_(2, 78)_ = 37.34, p < 0.001, *η*^²^_*p*_ = 0.49) and “pressure” (F_(2, 78)_ = 8.45, p < 0.001, *η*^²^_*p*_ = 0.18), indicating that paresthesia depended on duration as well as on intensity of the stimulus (but to a lesser extent:*η*^²^_*p*_ = 0.49 vs *η*^²^_*p*_ = 0.18). Post-hoc comparisons revealed statistically significant differences between all three durations (90s vs. 120s, 90s vs. 150s, 120s vs. 150s), however, in case of stimulus intensity (pressure), differences were significant only between 100 vs 150 mmHg and 100 vs 200 mmHg, but not between 150 mmHg vs 200 mmHg. No interaction of “pressure” × “time” factors was observed (F_(4, 156)_=2.37, p=0.55,). The stimulus-response function parameters are illustrated in **Figure 2**.

**Figure 2.**
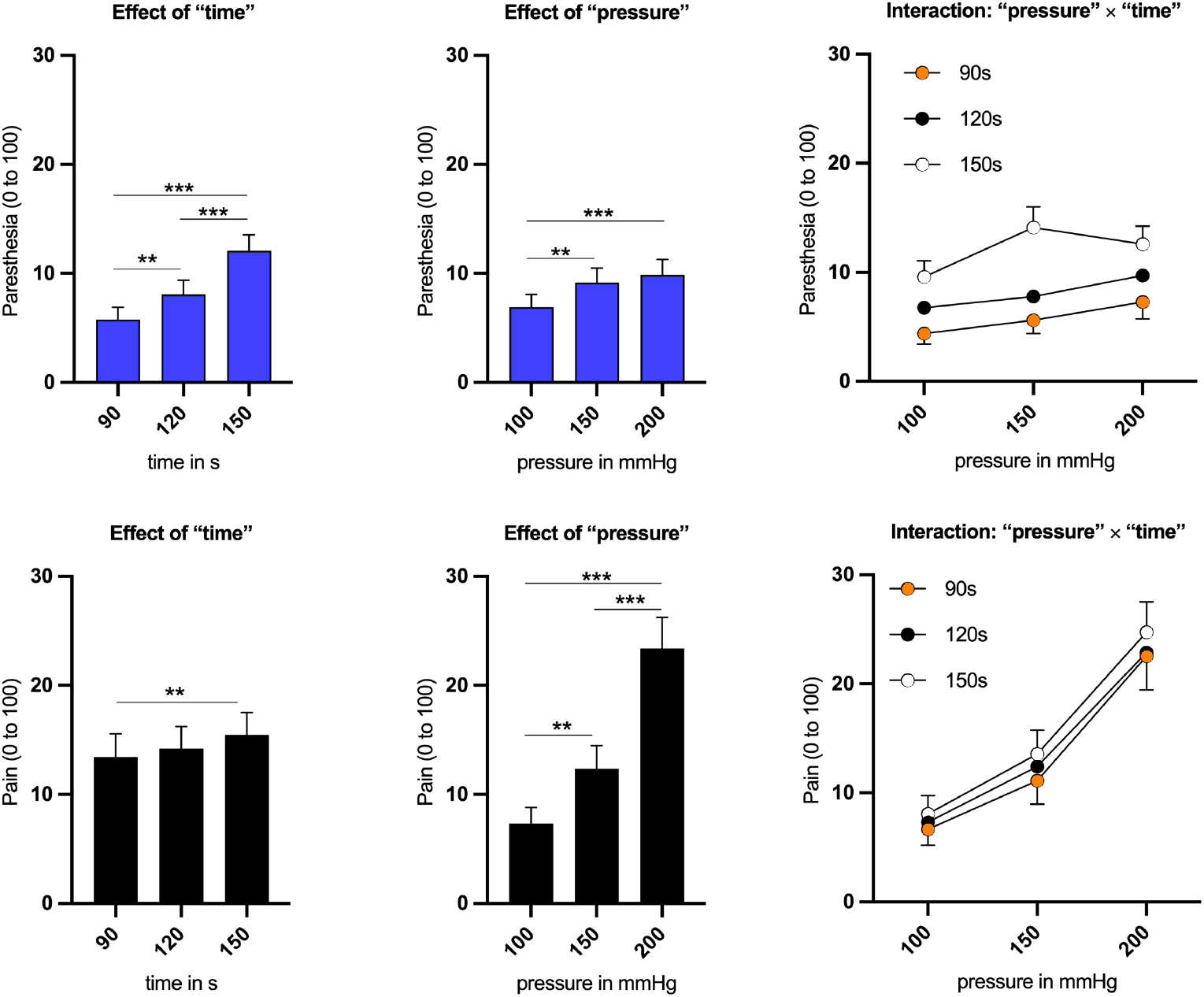
Main and interaction effects. The upper panel shows the average values of perceived paresthesia for each stimulus duration (upper left) and intensity (upper middle) and interaction effects (upper right) (p = 0.054). Steven’s power law function fitted to the data show exponent >1 for the relationship with duration (1.52) and exponent < 1 for the relationship with pressure (0.50) indicating an exponential and logarithmic relationship, respectively. Reversed relationships were found for pain (lower panel): exponential was the relationship with pressure (1.86) and the logarithmic saturation with duration (0.27). Legend: CoVAS – Computerized Visual Analogue Scale; mmHg – millimetres of mercury.

### 3.2. Pain

The GLM analysis for pain also revealed statistically significant main effects for the factors “time” (F_(2, 78)_ = 7.28, p < 0.05,*η*^²^_*p*_)) and “pressure” (F_(2, 78)_ = 63.02, p < 0.05, *η*^²^_*p*_ = 0.62). But contrary to paresthesia, *post-hoc* comparisons revealed statistically significant differences between all pressure intensities, however, in the temporal domain, significant differences were found only between the shortest and the longest time duration (90 vs. 150s). No interaction of “pressure” × “time” factors was observed (F_(4, 156)_=0.37, p=0.83, *η*^²^_*p*_ = 0.009 (**Figure 2**).

### 3.3. Electrodermal activity

The GLM analysis applied to SC data did not show statistically significant effects for “time” (F_(2, 58)_ = 0.86, p = 0.43, *η*^²^_*p*_ = 0.03), “pressure” (F_(2, 58)_ = 0.99, p = 0.38, *η*^²^_*p*_ = 0.03) and “time” × “pressure” interaction (F_(4,116)_ = 0.31, p = 0.87, *η*^²^_*p*_ *=* 0.01).

### 3.4. Between-session reliability of symptoms induction for average outcome

Intraclass correlation coefficients with 95% confidence intervals (CI), standard error of measurements (SEM) as well as smallest detectable differences of intensity of symptoms (pain, paresthesia) and signal (SC) are presented in **Table 3**. The overall agreement between first and second session is demonstrated on **Figure 3**. Averaged data from the first session plotted against data from the second session are shown as Bland-Altman plots. In general, negligible bias was found in the agreement for pain (2.54 ± 5.74) and paresthesia (−0.51 ± 3.93), and for both symptoms, 95% of the observations fall within the Limits of Agreement (± 1.96 SD of the absolute difference) [5].

**Table 3.** Reliability of pain, paresthesia, and skin conductance.

| Duration<br>[s] | Intensity<br>[mmHg] | Paresthesia |  |  |  | Pain |  |  |  | EDA |  |  |  |
| --- | --- | --- | --- | --- | --- | --- | --- | --- | --- | --- | --- | --- | --- |
|  |  | ICC<sub>(3,1)</sub> | 95% CI | SEM | SDD | ICC<sub>(3,1)</sub> | 95% CI | SEM | SDD | ICC<sub>(3,1)</sub> | 95% CI | SEM | SDD |
| 90 | 100 | 0.78 | 0.63 - 0.88 | 2.99 | 8.29 | 0.76 | 0.57 - 0.86 | 4.89 | 13.56 | 0.66 | 0.14 - 0.86 | 1.03 | 2.85 |
|  | 150 | 0.80 | 0.65 - 0.89 | 3.58 | 9.92 | 0.90 | 0.81 - 0.95 | 4.41 | 12.22 | 0.52 | 0.19 - 0.74 | 1.40 | 3.87 |
|  | 200 | 0.83 | 0.70 - 0.91 | 4.19 | 11.63 | 0.78 | 0.61 - 0.88 | 9.77 | 27.07 | 0.73 | 0.34 - 0.88 | 1.11 | 3.06 |
| 120 | 100 | 0.66 | 0.45 - 0.81 | 5.6 | 15.53 | 0.70 | 0.49 - 0.83 | 5.38 | 14.92 | 0.61 | 0.20 - 0.82 | 1.09 | 3.01 |
|  | 150 | 0.69 | 0.48 - 0.82 | 5.21 | 14.45 | 0.80 | 0.65 - 0.89 | 6.35 | 17.59 | 0.59 | 0.24 - 0.79 | 1.16 | 3.21 |
|  | 200 | 0.56 | 0.31 - 0.74 | 6.9 | 19.12 | 0.79 | 0.63 - 0.88 | 8.77 | 24.32 | 0.26 | -0.04 - 0.53 | 1.79 | 4.95 |
| 150 | 100 | 0.57 | 0.32 - 0.75 | 6.79 | 18.82 | 0.71 | 0.51 - 0.83 | 6.24 | 17.30 | 0.42 | 0.09 - 0.66 | 1.50 | 4.15 |
|  | 150 | 0.52 | 0.24 - 0.71 | 9.56 | 26.51 | 0.75 | 0.53 - 0.87 | 7.28 | 20.17 | 0.21 | -0.08 - 0.49 | 1.89 | 5.23 |
|  | 200 | 0.68 | 0.47 - 0.82 | 6.43 | 17.84 | 0.80 | 0.66 - 0.89 | 8.17 | 22.64 | 0.30 | -0.01 - 0.57 | 1.83 | 5.07 |

**Legend:** Note: Intraclass correlation coefficients (ICC, model 3,1) are presented with standard error of the measurement (SEM) and smallest detectable difference (SDD). Reliability is presented for three measured outcomes: pain intensity (Computerized Visual Analogue Scale, CoVAS), paresthesia intensity (CoVAS), electrodermal activity (EDA, physiological signal expressed in mS).

**Figure 3.**
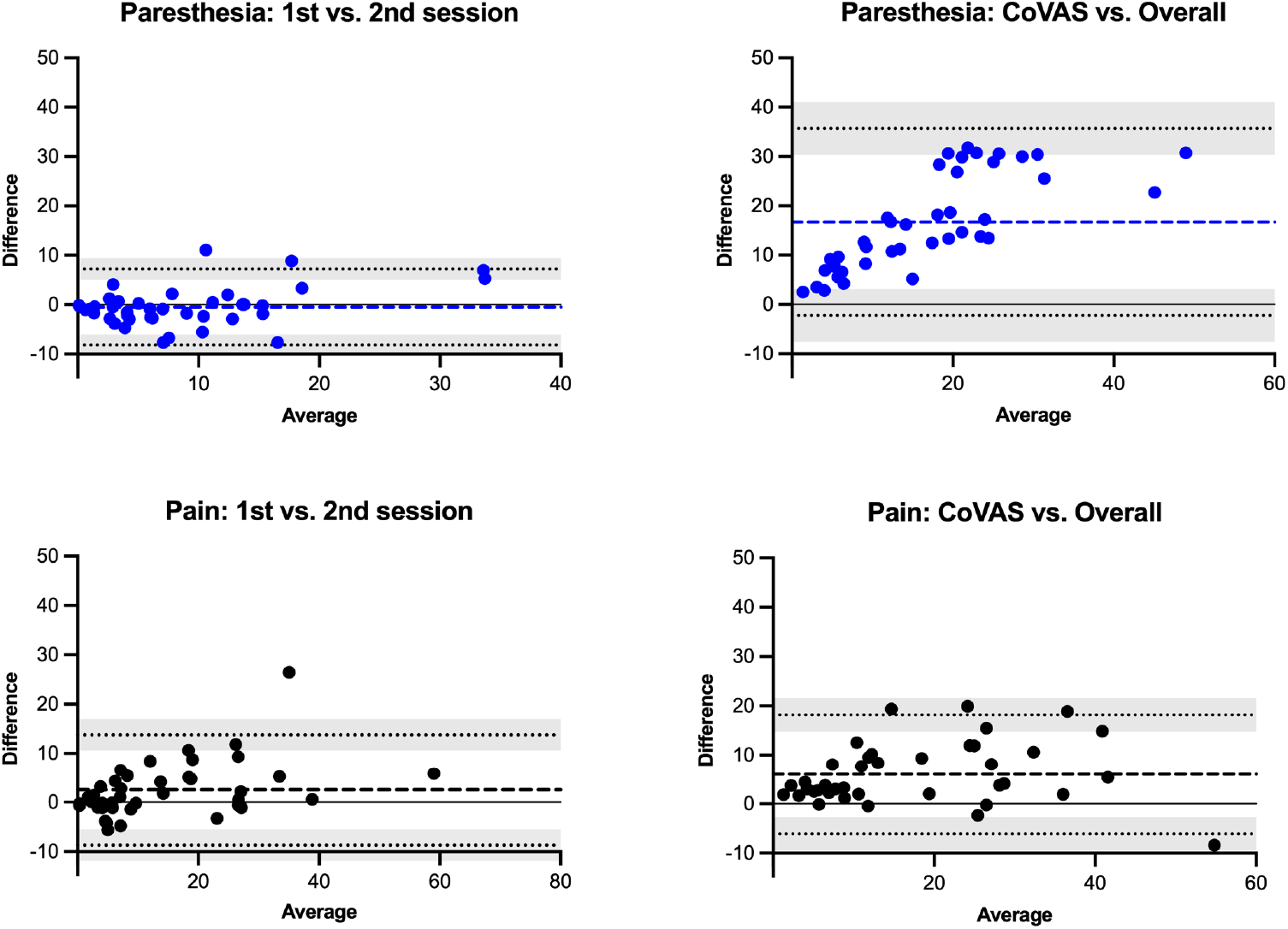
Bland-Altman plots. Agreement between first and second session is shown on the left and agreement between real-time and overall ratings on the right. Bland-Altman plots demonstrate individual differences (dots) between two ratings (first vs. second session on the left, or overall vs. CoVAS on the right) plotted against the average of ratings from two measures (methods). The thick horizontal line (paresthesia in blue, pain in black) illustrates the mean difference (bias) with its 95% limits of agreement (dotted line). The shaded area indicates the 95% CI to the upper and lower limits of agreement. The black horizontal line (zero) indicates absolute agreement.

### 3.5. Intraclass Correlation Coefficient

ICCs for paresthesia are characterized by generally higher reliability in the shortest stimul duration regardless of pressure intensity. Reliability was good (ICC_(3,1)_ = 0.78 - 0.83) for 90 seconds stimuli durations and decreased for longer durations, however, still obtaining moderate reliability (ICC_(3,1)_ = 0.51 - 0.69). Mostly good reliability for pain (ICC_(3,1)_ = 0.75-0.90) was observed in 7 out of 9 stimuli and moderate (ICC_(3,1)_ = 0.70-0.71) in the two remaining stimuli. ICCs for SC reached poor (ICC_(3,1)_ = 0.21 – 0.43) to moderate reliability (ICC_(3,1)_ = 0.52 – 0.73) (see **Table 3)**.

Similar analyses were performed based on AUC and comparable ICC values were obtained (see **Appendix S5**).

### 3.6. Relationship between real-time vs. overall ratings

Validity analyses revealed a significant correlation between real-time CoVAS ratings and (single) overall ratings collected after each trial by using a traditional VAS scale (**Table 4**). In general, agreement between the two methods to assess symptoms was acceptable for paresthesia with all observations falling within the 95% LoAs as indicated by Bland-Altman analysis (**Figure 3**). However, for both symptoms, bias was observed as indicated by a shift in mean differences between the two methods: overall ratings were higher than real-time for pain (diff. 6.03 ± 6.19) and paresthesia (difference 16.70 ± 9.68). Pearson correlations between the arm circumference and intensity ratings (pain and paresthesia) were not significant (paresthesia r = -0.09, p=0.56; pain r = -0.04, p=0.82; SC r = -0.20, p=0.21) (see **Appendix S6)**.

**Table 4.**
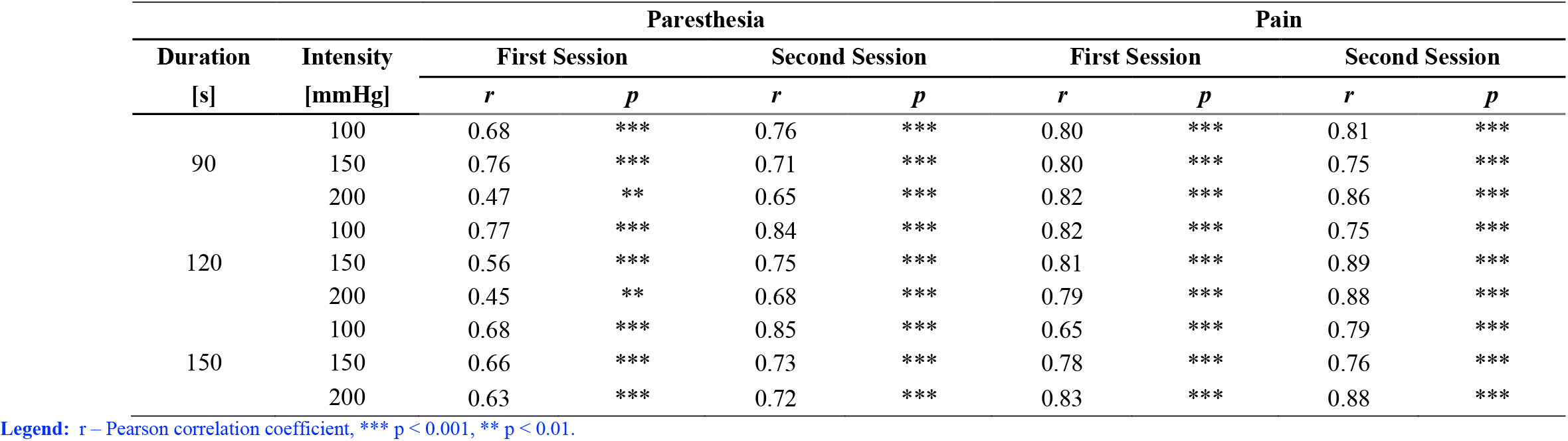
Results of correlations between CoVAS (real - time measurement) and overall ratings for paresthesia and pain.

## 4. DISCUSSION

This experiment systematically evaluated the effects of duration and intensity of noxious pressure stimuli on pain and paresthesia in young and healthy individuals using computer-controlled cuff algometry. The findings revealed distinct stimulus-response relationships: pain increased exponentially with pressure but was less influenced by duration, while paresthesia was primarily shaped by the duration of exposure and showed no significant increase with higher pressures. Pain remained relatively stable over time, whereas paresthesia developed more gradually. Reported symptoms demonstrated good reliability, in contrast to SC data, which showed poor reliability and no responsiveness to the stimulation. The different trajectories of pain and paresthesia development, as shown in Figure 1, along with their distinct psychophysical effects, demonstrate that the applied stimulation consistently and reliably induced these symptoms.

Neuropathic pain and paresthesia often result from damage or dysfunction within the nervous system [8,37] driven by a combination of peripheral factors (e.g., ion channel modulation, inflammatory mediators) [29] and central mechanisms (e.g., enhanced synaptic transmission, neuroplastic changes, loss of inhibitory control) [50]. Hypoxia and oxidative stress further contribute by triggering pro-inflammatory cytokines and reactive oxygen species (ROS), which sensitize nociceptors and intensify pain signaling [59]. In this study, cuff compression impacted multiple tissues, including skin, muscle, fascia, periosteum, and nerves. This compression activated mechanoreceptors, eliciting pain [43] while paresthesia resulted from reduced oxygen and nutrient delivery caused by prolonged compression and ischemia [54].

The results of this experiment demonstrated that paresthesia sensation primarily depended on the duration of compression rather than the stimulus intensity (pressure). As noted by Mogyoros et al. [38], paresthesia during prolonged ischemia is linked to increased axonal excitability, which begins within 30 seconds and plateaus within 3–5 minutes. In this study, ischemic stimuli lasted 90, 120, and 150 seconds, remaining within the window of increasing excitability. Each stimulus effectively induced ischemia, mimicking conditions typical for NP. Moreover, the intensity of paresthesia clearly saturated, as the additional step from 150 to 200 mmHg did not further increase symptoms, thereby confirming a nonlinear stimulus-response relationship. This finding may have clinical relevance, suggesting that stronger nociception does not necessarily exacerbate paresthesia.

A divergent pattern was observed for pain perception: pain increased exponentially with higher pressure, consistent with studies using this experimental pain model and with stimulus-response characteristics for mechanical [18], electrical [15], thermal [1] stimuli. Polianskis et al. [44] found a significant correlation (rs = 0.7) between pain and applied pressure to the calf muscles. Interestingly, pain intensity was not dependent on the duration of the stimulus as was the case for paresthesia. In a later study by Polianskis et al. [45], pain assessed over 10 minutes of compression remained stable for about two minutes after an initial increase in lower-intensity conditions, while moderate and severe pain showed adaptation after ∼140 seconds. Although further research must confirm this, it is likely that this sudden adaptation could be related to an interaction between pain and paresthesia that reached its peak after 150 seconds.

Given the simultaneous occurrence of pain and paresthesia in clinical settings, an experimental model was developed to induce and assess both symptoms concurrently. This study utilized computer-controlled cuff algometry for several key reasons. First, it is a widely used technique to induce pain [12,17,18,23,27,32,44], known for its good reliability [18,30], a finding also confirmed in the present study (ICC = 0.70 to 0.90). Second, compared to handheld algometry, computer-controlled cuff algometry allows for suprathreshold stimulation with varied temporal and intensity characteristics [21,22], crucial for inducing not only pain but also paresthesia, which requires prolonged oxygen restriction [54]. Tonic compression was therefore deemed the most suitable approach.

Additionally, this method facilitates real-time monitoring of sensory changes during stimulation via CoVAS, a feature essential to this experiment [18,44]. Real-time monitoring offers several significant advantages in the assessment of NP and paresthesia. By providing continuous, immediate feedback on sensory changes, this approach enhanced both the accuracy and relevance of the data collected.

Real-time assessment captures dynamic changes in pain and paresthesia, offering more precise and accurate measurements compared to retrospective self-reports by minimizing recall bias and accurately recording transient symptoms [51]. Interestingly, although a significant correlation was found between overall pain ratings and CoVAS ratings, overall ratings tended to be higher, particularly for paresthesia. This discrepancy likely reflects paresthesia’s gradual development compared to pain’s more stable trajectory. Consequently, recalling overall pain ratings appeared less cognitively demanding than paresthesia ratings. Furthermore, the “peak-end” artifact—where symptoms are rated based on peak intensity and stimulus endpoint—may have had a greater influence on paresthesia ratings than on pain [24].

Surprisingly, no effect of stimulus parameters was found on SC, nor was any correlation observed between SC signals and paresthesia or pain. However, the literature on SC and pain is inconsistent, with some studies showing a relationship while others do not. In some studies, authors have argued that SC can encode the intensity of a stimulus rather than the intensity of pain [33,40]. Additionally, others, such as Scheuren et al. [48], suggest that SC may be influenced by factors beyond just stimulus-associated arousal or subjective pain intensity. Meanwhile, other research has found that SC responses can discriminate between different levels of reported pain [7,16,36]. In contrast, Breimhorst et al. [7], demonstrated that SC responses can distinguish between different levels of pain appraisal, though their use of phasic rather than tonic stimulation may explain the differing results. In this study, SC was measured on the same side as the nociceptive stimulation, and prolonged compression could have impaired sympathetic nerve transduction, introducing variability. Additionally, the relatively low pain levels reported may have caused a floor effect, as SC responses are typically more pronounced with higher pain intensity [36]. The maximal stimulus of 250 mmHg for 150 seconds was chosen to ensure safety and participant tolerance, potentially limiting SC responsiveness. Future research should investigate SC responses in limb ischemia and alternative stimulation protocols, as the literature in this area remains limited.

The findings of this current study have significant implications for experimental pain research, particularly in advancing the understanding of NP conditions, while also offering insights relevant to clinical practice: The ability to simultaneously measure pain and paresthesia using computer-based cuff algometry offers a more nuanced perspective on these symptoms and their interrelationship, enhancing diagnostic and therapeutic approaches. NP is complex and often involves co-occurring symptoms such as pain and paresthesia. Traditional pain models typically focus on a single symptom, missing the intricate interactions between these symptoms. By reliably inducing and measuring pain and paresthesia, this study provides a more comprehensive surrogate model for investigating neuropathic pain.

This dual-symptom approach offers several benefits. First, it enables researchers to dissect mechanistic pathways, shedding light on the distinct and overlapping neural mechanisms underlying NP. Second, it allows for the characterization of symptom interactions, capturing the temporal dynamics and intensity-response relationships to better understand how these symptoms influence each other and the overall pain experience.

Clinically, this method holds promise for more accurate monitoring of treatment efficacy and recovery. Patients with entrapment neuropathies, such as cervical radiculopathy, commonly experience arm pain accompanied by paresthesia [52]. Tampin and colleagues found that these patients exhibit functional impairments in their maximal pain area (e.g., the arm), including reduced mechanical detection thresholds and lower pressure pain sensitivity on the symptomatic side compared to the asymptomatic side or healthy controls. This model could support patient outcome monitoring, as nerve healing and function restoration would likely lead to increased pain levels during stimulation relative to baseline, eventually aligning with the asymptomatic side over time.

Additionally, evaluating how interventions affect both, pain and paresthesia, provides a holistic measure of therapeutic success, extending beyond pain reduction, alone. Since these two symptoms often co-occur in clinical practice, assessing only one of them may limit our understanding of how interventions targeting one symptom could influence the other. By simultaneously assessing both symptoms, this method offers a unique opportunity to examine interdependencies between pain and paresthesia. For instance, an improvement in one symptom might lead to alleviation of the other, potentially mediated by generalization of specific or non-specific treatment effects [9,55,58].

### Limitations and future directions

This study has several limitations. Firstly, the lack of racial and ethnic diversity in the sample is a constraint, as all participants identified as white, reflecting the demographics of the study region in Poland. This limited access to participants from more diverse backgrounds may restrict the generalizability of the findings. Future studies should aim for a more diverse participant pool to enhance external validity.

Secondly, the definition of “healthy, pain-free” participants was broad. While individuals with known medical conditions or pain-related disorders were excluded, we did not use the detailed assessment criteria recommended by EUROPAIN and NEUROPAIN consortia. This may have overlooked subtle baseline health differences, limiting control over these factors. Future studies should adopt stricter screening protocols to improve participant characterization and reduce variability.

A third limitation is the lack of systematic assessment of participants’ ability to rate pain and paresthesia simultaneously. While verbal feedback suggested no major difficulties, the absence of a formal evaluation, such as an exit questionnaire, limits understanding of potential challenges. Future studies should include structured assessments to validate this dual-rating approach.

Lastly, the applicability of this method to conditions with paresthesia not induced by compression, such as fibromyalgia or neuropathic pain syndromes, is uncertain. These conditions may involve distinct mechanisms, potentially limiting the model’s relevance. Validation in such populations is needed to assess reliability and determine whether adaptations are required.

## 5. ACKNOWLEGDEMENTS

The study took place in the certified Laboratory of Pain Research (ISO 9001:2015). The study was financed by Polish National Science Center grant number (2020/37/B/HS6/04210).

