## Appendix for "Experimentally induced pain and paresthesia respond differently to parameter changes of cuff-based compression in pain-free young individuals"

### Appendix (supplementary materials)

#### S1. Means and SDs for all conditions and outcomes for AUC (*$10^{-3}$)

| Condition | Paresthesia | Pain | EDA |
|---|---|---|---|
| (n=40) | mean (SD) | mean (SD) | mean (SD) |
| 90/100 | 0.40 (0.55) | 0.60 (0.84) | 50.43 (29.88) |
| 90/150 | 0.50 (0.69) | 1.00 (1.23) | 49.17 (31.00) |
| 90/200 | 0.66 (0.88) | 2.04 (1.77) | 51.36 (35.19) |
| 120/100 | 0.81 (1.06) | 0.88 (1.09) | 64.17 (36.97) |
| 120/150 | 0.93 (1.03) | 1.50 (1.63) | 64.48 (38.10) |
| 120/200 | 1.16 (1.11) | 2.75 (2.17) | 64.47 (38.35) |
| 150/100 | 1.44 (1.38) | 1.21 (1.61) | 77.34 (47.31) |
| 150/150 | 2.11 (1.80) | 2.04 (2.05) | 78.89 (47.84) |
| 150/200 | 1.88 (1.57) | 3.72 (2.65) | 81.66 (50.63) |

**Legend:** Condition i.e., 90/100 indicates condition where the pressure stimulus had a duration of 90 seconds and intensity of 100 mmHg; auc – area under the curve; SD – Standard Deviation; EDA- electrodermal activity; AUC values are reduced and presented as value *$10^{-3}$

#### S2. Results of General Linear Model (GLM) analysis conducted on paresthesia symptoms (Area Under the Curve)

| Main or interaction effect | df | F | p |
|---|---|---|---|
| "session" | 1, 39 | 0.74 | 0.40 |
| "time" | 2, 78 | 61.66 | **< 0.001** |
| "pressure" | 2, 78 | 7.66 | **< 0.001** |
| "session" × "time" | 2, 78 | 0.50 | 0.61 |
| "session" × "pressure" | 2, 78 | 0.84 | 0.44 |
| "time" × "pressure" | 4, 156 | 3.00 | **0.02** |
| "session" × "time" × "pressure" | 4, 156 | 0.85 | 0.50 |

Note: The table above presents main and interaction effects of the GLM analysis. Bold values indicate statistically significant effect. Post-hoc comparisons for significant 2-way interaction ($F_{(4,156)} = 3.00$, $p = 0.02$, $\eta^2_p=0{,}07$) are presented below:

| Intensity / pressure | $150_{mmHg}$ | $200_{mmHg}$ | $150_{mmHg}$ | $200_{mmHg}$ | $150_{mmHg}$ | $200_{mmHg}$ |
|---|---|---|---|---|---|---|
| Duration: | 90s | | 120s | | 150s | |
| **$100_{mmHg}$** | 1.00 | 0.63 | 0.99 | 0.20 | **< 0.001** | **0.04** |
| **$150_{mmHg}$** | | 0.98 | | 0.77 | | 0.77 |

Note: Given that the area under the curve (AUC) is time-dependent i.e. it increases with the duration of the stimulus, no valid inference is possible from comparisons of the same stimulus intensity across different stimulus durations. Therefore, only comparisons between different intensities (pressures) within the same duration domain are reported below.

**S3. Results of General Linear Model (GLM) analysis conducted on pain (Area Under the Curve)**

| Main or interaction effect | df | F | p |
|---|---|---|---|
| "session" | 1, 39 | 7,52 | **< 0.01** |
| "time" | 2, 78 | 56.67 | **< 0.001** |
| "pressure" | 2, 78 | 64.18 | **< 0.001** |
| "session" × "time" | 2, 78 | 0.36 | 0.70 |
| "session" × "pressure" | 2, 78 | 0.86 | 0.43 |
| "time" × "pressure" | 4, 156 | 11.48 | **< 0.001** |
| "session" × "time" × "pressure" | 4, 156 | 1.58 | 0.18 |

Note: The table above presents main and interaction effects of the GLM analysis. Bold values indicate statistically significant effect. Post-hoc comparisons for significant 2-way interaction ($F_{(4,156)} = 11.48$, $p < 0.001$, $\eta^2_p = 0.23$) are presented below:

| **Intensity / pressure** | **$150_{mmHg}$** | **$200_{mmHg}$** | **$150_{mmHg}$** | **$200_{mmHg}$** | **$150_{mmHg}$** | **$200_{mmHg}$** |
|---|---|---|---|---|---|---|
| Duration: | 90s | | 120s | | 150s | |
| **$100_{mmHg}$** | **0.01** | **< 0.001** | **< 0.001** | **< 0.001** | **< 0.001** | **< 0.001** |
| **$150_{mmHg}$** | | **< 0.001** | | **< 0.001** | | **< 0.001** |

Note: Given that the area under the curve (AUC) is time-dependent i.e. it increases with the duration of the stimulus, no valid inference is possible from comparisons of the same stimulus intensity across different stimulus durations. Therefore, only comparisons between different intensities (pressures) within the same duration domain are reported below.

**S4. Results of General Linear Model (GLM) analysis conducted on electrodermal activity (Area Under the Curve)**

| Main or interaction effect | df | F | p |
|---|---|---|---|
| "session" | 1, 33 | 13.44 | **< 0.001** |
| "time" | 2, 66 | 47.37 | **< 0.001** |
| "pressure" | 2, 66 | 1.05 | 0.36 |
| "session" × "time" | 2, 66 | 3.46 | **0.04** |
| "session" × "pressure" | 2, 66 | 0.91 | 0.41 |
| "time" × "pressure" | 4, 132 | 0.16 | 0.63 |
| "session" × "time" × "pressure" | 4, 132 | 0.62 | 0.65 |

Note: The table above presents main and interaction effects of the GLM analysis. Bold values indicate statistically significant effect. No significant 2-way interaction was found.

**S5. Reliability of measurement reported for pain, paresthesia, and skin conductance (Area Under the Curve)**

| Duration [s] | Intensity [mmHg] | Paresthesia | | | | Pain | | | | EDA | | | |
|---|---|---|---|---|---|---|---|---|---|---|---|---|---|
| | | $ICC_{(3,1)}$ | 95% CI | SEM | SDD | $ICC_{(3,1)}$ | 95% CI | SEM | SDD | $ICC_{(3,1)}$ | 95% CI | SEM | SDD |
| | 100 | 0.78 | 0.63 - 0.88 | 2.69 | 7.47 | 0.76 | 0.57 - 0.86 | 4.42 | 12.25 | 0.67 | 0.15 - 0.86 | 190.42 | 527.82 |
| 90 | 150 | 0.80 | 0.65 - 0.89 | 3.22 | 8.92 | 0.90 | 0.81 - 0.95 | 3.99 | 11.05 | 0.52 | 0.18 - 0.74 | 255.57 | 708.42 |
| | 200 | 0.83 | 0.70 - 0.91 | 3.78 | 10.49 | 0.78 | 0.61 - 0.88 | 8.84 | 24.49 | 0.74 | 0.36 - 0.89 | 196.60 | 544.96 |
| | 100 | 0.67 | 0.45 - 0.81 | 6.71 | 18.61 | 0.70 | 0.49 - 0.83 | 6.48 | 17.96 | 0.63 | 0.23 - 0.82 | 254.45 | 705.30 |
| 120 | 150 | 0.69 | 0.48 - 0.82 | 6.27 | 17.37 | 0.80 | 0.65 - 0.89 | 7.64 | 21.19 | 0.58 | 0.23 - 0.78 | 280.93 | 778.69 |
| | 200 | 0.56 | 0.31 - 0.74 | 8.29 | 22.98 | 0.79 | 0.63 - 0.88 | 10.57 | 29.29 | 0.26 | -0.04 - 0.53 | 431.60 | 1196.32 |
| | 100 | 0.57 | 0.32 - 0.75 | 10.20 | 28.28 | 0.71 | 0.51 - 0.83 | 9.38 | 26.01 | 0.41 | 0.09 - 0.66 | 449.66 | 1246.38 |
| 150 | 150 | 0.51 | 0.24 - 0.71 | 14.39 | 39.88 | 0.75 | 0.53 - 0.87 | 10.93 | 30.31 | 0.21 | -0.08 - 0.49 | 565.99 | 1568.86 |
| | 200 | 0.68 | 0.47 - 0.82 | 9.67 | 26.81 | 0.80 | 0.66 - 0.89 | 12.28 | 34.05 | 0.30 | -0.01 - 0.56 | 548.30 | 1519.80 |

Note: Intraclass correlation coefficients (ICC, model 3,1) are presented with standard error of the measurement (SEM) and smallest detectable difference (SDD). Reliability is presented for three measured outcomes: pain intensity (Computerized Visual Analogue Scale, CoVAS), paresthesia intensity (CoVAS), electrodermal activity (EDA, physiological signal expressed in mS).

**S6. Results of the correlation between arm circumference and the average outcome derived from all conditions using the Pearson-product coefficient.**

| Outcome | Average | |
|---|---|---|
| | r | p |
| paresthesia | -0,09 | 0,56 |
| pain | -0,04 | 0,82 |
| EDA | -0,20 | 0,21 |

**S7. Discussion**

*Mechanisms and Interactions of Neuropathic Pain and Paresthesia*

Neuropathic pain is a complex, chronic pain state that is typically accompanied by tissue injury. The nerve fibers themselves may be damaged, dysfunctional, or injured. These damaged nerve fibers send incorrect signals to other pain centers. Neuropathic pain can be caused by a variety of conditions, including diabetes, shingles, multiple sclerosis, trigeminal neuralgia, and postherpetic neuralgia [6,12]. Neuropathic pain affects approximately 7-10% of the general population [6] and account for approximately 15-25% of all chronic pain cases [5]. It is more common among individuals with certain conditions such as diabetes, where it affects up to 26% of patients [6]. Paresthesia, characterized by abnormal sensations such as tingling, burning, or numbness, often accompanies neuropathic pain [15]. This symptom is frequently reported in conditions that involve nerve damage, including carpal tunnel syndrome [4], radicular pain from spinal nerve root compression [3], and peripheral neuropathy [2]. The impact of neuropathic pain and paresthesia on quality of life is profound. These symptoms can lead to significant physical, emotional, and psychological distress. Patients with neuropathic pain often experience severe discomfort, sleep disturbances, anxiety, and depression [13,14]. The chronic nature of the pain can also lead to difficulties in performing daily activities, which further exacerbates the psychological burden [7]. Neuropathic pain is notoriously difficult to treat. Conventional analgesics, such as NSAIDs and opioids, are often ineffective [8]. Treatment usually requires a combination of medications, including antidepressants, anticonvulsants, and topical agents, along with non-pharmacological approaches such as physical therapy and psychological support [1].

*Relationship between arm circumference and intensity of symptoms*

In the experiment, we additionally aimed to investigate the relationship between anthropometric data and intensity of symptoms. Pressure algometry activates receptors present in both superficial and deep tissues [11]. We expected that individuals with larger arm circumference, which indicates a greater volume of tissue and a higher number of potential receptors, may experience more intense symptoms compared to those with smaller arm circumference. In the study of Graven-Nielsen [9] for example, pressure pain thresholds were significantly higher when measured on lower limbs compering to upper limbs. In our study we found no correlation between arms circumferences and perceived pain. It may be due to several reasons. One of the potential explanations could be that the variations in arm girth among the subjects were too minimal to observe any significant differences in perceived symptoms (with SD of 4,06 cm) when compared to the differences observed between the leg and arm in the study. On the other hand, according to the study conducted by Jensen et al. [10], the subcutaneous administration of lidocaine as an analgesic led to a significant 70% increase in the pressure pain threshold (PPT) value. That indicates that the cutaneous and subcutaneous components contribute significantly greater to the perception of pain than underlying tissues.

*Reliability and validity*

Secondary aim was to provide preliminary reliability and validity data of the technique designated to measure two symptoms simultaneously. Inter-session reliability for pain was demonstrated to be mostly good (7/9 stimuli) indicating, that this method of inducing pain is reliable. It is in line with the study of Graven-Nielsen et al. [9] where good to excellent (0.60-0.90) interclass correlation were demonstrated for pressure pain threshold measured on the upper extremity using computer-based cuff algometry. On the other hand, paresthesia reached good reliability for shortest stimuli durations and the reliability decreased along with increasing stimulus duration, but still reaching moderate reliability. The above suggests that proposed model of concurrently induced pain and paresthesia might be reliable and could be used in experiments that requires the induction of two symptoms at the same time.

The knowledge derived from this experiment will allow to use novel experimental model that can be used to induce two symptoms simultaneously in a controlled manner, i.e., by adjusting the pressure parameters in such a way as to induce symptoms of the intended parameters.

### S8. Instruction and informations to participants

"There will be a device between your hands to assess pain and tingling. There are three sliders on it. The slider on the left is used to determine the value of the pain you feel, where 0 means no pain and 10 means the worst pain you can imagine. The middle slider is used to determine the value of the tingling sensation felt, where 0 means no tingling and 10 means the worst tingling you can imagine. The slider on the right is inactive and will not be used."

"Please try to operate the sliders with your non-examining hand and maneuver them continuously to reflect the current state of tingling or pain felt. The aim of the examination is to detect as little change as possible in the severity of your symptoms, so it is crucial that you assess these symptoms on an ongoing basis, during every second of the examination."

Explaining the concept of "fear of pain"
"Fear of pain is how worried or scared you feel about the possibility of experiencing pain. On a scale from 0 to 10, we'd like you to think about how much fear or concern you feel right now about the pain you might experience from the procedure. A **0** means you feel no fear at all—you're not worried or scared about the pain. A **10** means you feel the worst fear you can imagine—you're extremely worried or scared about the pain. Please choose the number that best matches how you feel right now."
